# No evidence that HLA genotype influences the driver mutations that occur in cancer patients

**DOI:** 10.1101/2021.04.21.440830

**Authors:** Noor Kherreh, Siobhán Cleary, Cathal Seoighe

**Affiliations:** School of Mathematics, Statistics and Applied Mathematics, National University of Ireland Galway, Galway, Ireland

**Keywords:** Cancer, Driver mutations, MHC, Adaptive immune response

## Abstract

The major histocompatibility (MHC) molecules are capable of presenting neoantigens resulting from somatic mutations on cell surfaces, potentially directing immune responses against cancer. This led to the hypothesis that cancer driver mutations may occur in gaps in the capacity to present neoantigens that are dependent on MHC genotype. If this is correct, it has important implications for understanding oncogenesis and may help to predict driver mutations based on genotype data. In support of this hypothesis, it has been reported that driver mutations that occur frequently tend to be poorly presented by common MHC alleles and that the capacity of a patient’s MHC alleles to present the resulting neoantigens is predictive of the driver mutations that are observed in their tumour. Here we show that these reports of a strong relationship between driver mutation occurrence and patient MHC alleles are a consequence of unjustified statistical assumptions. Our reanalysis of the data provides no evidence of an effect of MHC genotype on the oncogenic mutation landscape.

## 1 Introduction

The immune system has evolved to recognize aberrant and non-self molecules, resulting from pathogen infection, somatic mutations and malformed proteins. The major histocompatibility complex (MHC) plays a key role in this process. There are two classes of MHC molecules, class I (MHC-I) and class II (MHC-II), encoded, in human, by a cluster of genes on chromosome 6. The human MHC genes and proteins, which are often termed Human Leukocyte Antigens (HLA), are diverse, with over 15,000 alleles identified [1]. Somatic mutations in genes encoding self proteins can result in an altered amino acid sequence, thereby generating so-called neo-antigens that have the potential to elicit an immune response upon presentation by the MHC to T-cells [2]. Through the killing of cells carrying immunogenic neoantigens, the immune system has been proposed to a play key role in shaping the cancer genome in a process referred to as immuno-editing [3, 4].

Dunn et al. first proposed the term immuno-editing to describe the dual ability of the immune system to defend the host by suppressing tumour growth and to shape the immunogenicity of tumours [5]. It is characterised by three phases – elimination, equilibrium and escape, collectively termed the three Es of cancer immuno-editing [5, 6]. The elimination phase involves the recognition and destruction of tumour cells by the immune system, before it is clinically detectable. Some cells are thought to escape elimination and enter into the equilibrium phase during which the immune system keeps tumour growth in check but cannot fully eliminate it. The tumour may continue to develop mutations that enable it to evade immune responses, resulting in a population of cells that are resistant to the immune response [6, 4]. The final stage occurs when the cancer escapes immune control, leading to uncontrolled proliferation, due potentially to reduced immunogenicity of cancer cells or to mutations that create an immunosuppressive environment [6, 3].

One of the mechanisms through which cancer evades the immune response is to acquire mutations that alter antigen presentation [7]. The most selective step of the process of antigen presentation to the immune cells is the binding of antigenic peptides to the MHC. This has been inferred by a variety of studies of the implications of mutating the HLA genes or the B2M gene, whose product, *β*_2_m, forms an integral part of MHC Class I molecules [8–12]. Loss or mutation of HLA or B2M genes is associated with an increase in tumour mutation burden [12]. A lack of neoantigens capable of eliciting an immune response could also allow cancers to avoid immune responses and several studies have reported selection against immunogenic somatic mutations in cancer [13, 9, 14], though the evidence for depletion of mutations that give rise to neoantigens has recently been questioned [15].

Here, we reanalyzed the data from two high-profile studies [16, 17] that reported that the driver mutations that are found in cancer patients can be predicted from the capacity of the patient’s MHC molecules to bind the resulting neoantigens. The patient harmonic mean best rank (PHBR) score was proposed in [16] and [17] as a measure of whether a neoantigen resulting from a somatic mutation can be bound by MHC molecules, given the HLA genotype of a patient. The score is derived from predicted binding affinities of the patient’s MHC molecules for the peptides spanning the mutation. The conclusions of both studies are based on an analysis of 1,018 cancer driver mutations in patients from the cancer genome atlas (TCGA). The focus of the 2017 study is on MHC class I alleles and the primary focus of the 2018 study is on presentation of cancer neoantigens by MHC class II molecules. The data for both comprised a binary matrix of mutation occurrences (indicating whether the driver mutation in each column has been observed in the patient in each row) and a matrix of PHBR scores corresponding to 9,176 and 5,942 patients for MHC class I and class II alleles, respectively. We reanalyzed these data and found that the conclusion of both papers, that cancer driver mutations emerge preferentially in gaps in the patient’s capacity to present neoantigens on MHC molecules, are not robust. We found that there is no evidence from the data that the driver mutations seen in a patient are influenced by the patient’s MHC class I or class II genotypes.

## 2 Methods

### Data

We performed a reanalysis of cancer driver mutations in TCGA and their predicted immunogenicities, reported in [16] and [17]. Both papers calculate a score that is used to predict the extent to which neoantigens are presented on MHC-I or MHC-II molecules, given the patient genotype. The score is calculated by considering all peptides of a specific length or range of lengths that contain the mutation. A rank-based presentation score was obtained for each peptide using NetMHC-pan3.0 [18] and for each of the patient’s HLA alleles the best rank value was retained. The PHBR score is then the harmonic mean (across the patient’s HLA alleles) of these best-rank scores (see [16] and [17] for details). This score was calculated for class I MHC alleles in [16] where it was based on peptides with lengths ranging from 8 – 11 amino acids and for class II alleles in [17], where it was based on peptides of length 15 amino acids. We applied the methodology as described to the TCGA data to obtain a binary matrix of driver mutation occurrences across patients and matrices of PHBR-I and PHBR-II scores across patients for each driver mutation. In order to ensure our results were precisely comparable to the published results we also requested the data matrices that were the basis of the original studies and these were kindly provided by the authors (following confirmation of the appropriate data access permissions).

### Logistic regression models relating mutation occurrences to PHBR scores

Following the notation of [16], consider a mutation matrix, with entries *y*_*ij*_ ∈ {0, 1}, indicating the presence or absence of driver mutation *j* in patient *i* and a matrix of PHBR-I or PHBR-II scores with real-valued entries, *x*_*ij*_, corresponding to the score of mutation *j*, given the MHC alleles of individual *i*. Two mixed effects logistic regression models were used in [16] to relate the log-odds of *y*_*ij*_ = 1 to the log of *x*_*ij*_. The first model, referred to as the *within-mutation* model, has a normally-distributed random effect,*β*_*j*_, that models differences in the frequencies of different driver mutations:

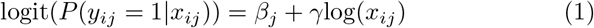

The second model, referred to as the *within-patient* model, uses a random effect, *η*_*i*_, to model differences in the abundance of driver mutations between patients, but does not model differences in the frequencies with which different driver mutations occur:

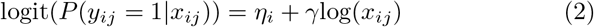

### Simulation

We designed a simple simulation scenario to illustrate how spurious results can be obtained from the within-patient model due to a failure to account for non-independence of the PHBR scores across patients (some driver mutations tend to have higher scores across patients, while others have lower scores, leading to the high degree of correlation in the scores of driver mutations between patients seen in Fig. 1A). The simulation consisted of 100 driver mutations, one of which had a high frequency (20% of 500 patients) and a relatively high PHBR score (normally distributed across patients with mean 10 and standard deviation 2). The remaining mutations occurred at low frequency (1%) and had normally distributed PHBR scores with mean 5 and standard deviation 2. We then fitted the within-patient model to this simulated dataset.

**Fig. 1.**
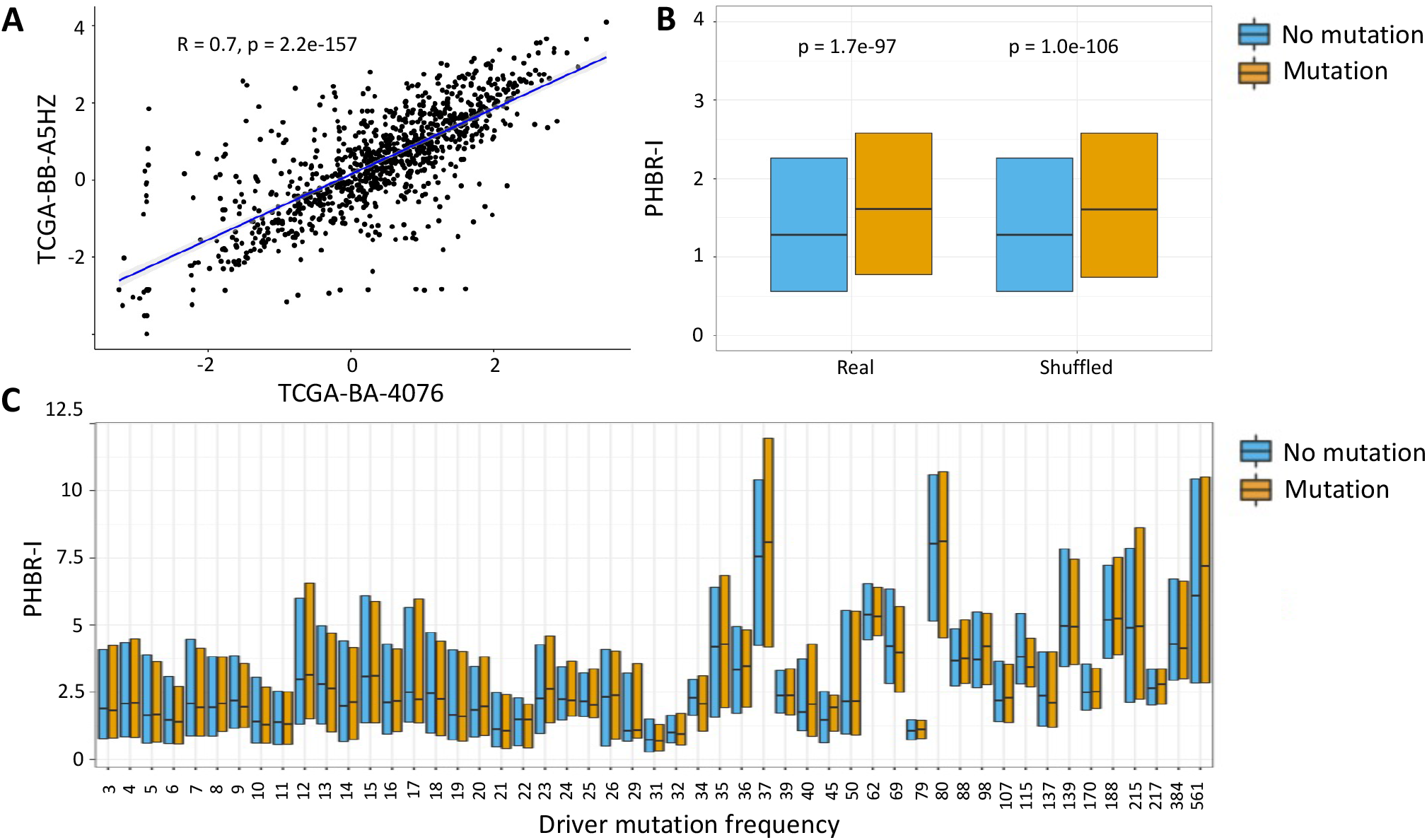
(A) Scatterplot of log PHBR-I scores of all driver mutations, calculated using the HLA genotypes of two randomly selected patients from TCGA. (B) Median and interquartile range of PHBR-I score in the No Mutation (blue) and Mutation (orange) groups for the real data and for data in which the MHC genotypes have been randomized between patients. (C) Median and interquartile range of PHBR-I scores in the No Mutation (blue) and Mutation (orange) groups in bins of mutation recurrence.

### Relationship between MHC-I coverage and cancer risk in UK Biobank

We retrieved HLA class I alleles from participants in the UK Biobank. These alleles were inferred using HLA*IMP:02 [19]. Only alleles that were called with imputation posterior probability greater than 0.5 and only participants with six HLA class I alleles called were retained. This left a total of 377,790 individuals. For each individual we determined the driver mutation coverage as the number of driver mutations with PHBR-I scores < 2, given the individual’s HLA genotype. We retrieved the self-reported cancer status (data field 20001) for these individuals. Treating the self-report of any cancer type as a case, we fitted a logistic regression model to case status as a function of age, sex and PHBR-I coverage.

## 3 Results

Using the predicted immunogenicities of driver mutations derived by [16] and [17], we re-investigated the relationship between immunogenicity and driver mutation occurrence across patients. In both [16] and [17] the predicted capacity of the MHC to present cancer driver mutations was compared between patients with and without the mutation. Higher values of the PHBR score (corresponding to low predicted capacity to bind neoantigens resulting from the mutation) in the patients in which the driver mutations occur was presented as evidence that driver mutations preferentially arise in patients who lack the MHC alleles that are capable of presenting them to T cells. In these comparisons of groups of PHBR scores, one group consists of the scores of driver mutations in patients in which the mutation is present (the Mutation group) and the other group (the No Mutation group) consists of PHBR scores of the driver mutations in the patients without the mutation. A given driver mutation can appear many times in the Mutation group in these comparisons -once for each patient in which it occurs. This is problematic, because the PHBR scores of mutations are highly correlated (Fig. 1A) and, thus, the data points are not independent. For example, a driver mutation that occurs in 500 patients will contribute 500 PHBR scores to the Mutation group and N – 500 scores to the No Mutation group, where N is the total number of patients. If the PHBR score of the mutation is generally high or generally low across patients, it will clearly have a disproportionate impact on the distribution of PHBR scores in the Mutation group.

The correlation in PHBR scores between patients is not solely due to sharing of HLA alleles. Even the PHBR scored using HLA alleles from different allele groups are significantly correlated (Fig. S1), but the scores of driver mutations were effectively treated as independent observations by the studies that reported an effect of HLA alleles on driver mutations. [17] used a statistical test (the Mann-Whitney U test) to compare the median PHBR-II score between the Mutation and No Mutation groups and reported a higher median score in the mutation group with a p-value < 2.2 × 10^*−*16^. This was interpreted as evidence that the patient HLA genotype influences the driver mutations that occur in cancer patients. However, the fundamental assumption of the test is that the observations in each group are independent and this assumption is clearly violated. We found that the differences between the Mutation and No Mutation groups are, in fact, just as large when the MHC genotypes are randomized between patients, indicating that this difference is not driven by patient genotype (Fig. 1B). Moreover, when we compared PHBR scores, grouped by driver mutation frequency (so that each driver mutation contributes the same number of observations to the Mutation group in each comparison), we saw no consistent differences (Fig. 1C).

In 100 randomizations of the HLA class I genotypes the median PHBR-I score of the Mutation group in the randomized data in fact exceeded the median of the Mutation group in the real data 94 times (p = 0.94 for the one-sided randomization-based test for a higher PHBR-I score in the Mutation group). Similarly, when we shuffled the HLA class II genotypes, the median PHBR-II score of the Mutation group in the shuffled data exceeded that of the real data 36 times (p = 0.34). Thus, comparison of PHBR scores between the Mutation and No Mutation group does not provide any support for the hypothesis that driver mutations occur preferentially in patients with MHC molecules that are not capable of binding the resulting neoantigens. In [16] and [17] PHBR scores of driver mutation occurrences were also compared against scores of occurrences for different mutation classes (e.g. germline mutations and passenger mutations). Because they contribute many times to the Mutation group, the existence of a small number of highly recurrent cancer driver mutations with high PHBR scores (i.e. low binding affinity) may be sufficient to skew all of these comparisons. This problem is compounded by the fact that the 1,018 driver mutations that are the basis of this study occur on just 168 different genes and PHBR scores are statistically significantly correlated between mutations in the same gene, particularly for class II alleles (Fig. S2). The number of distinct genes among the most highly recurrent cancer driver mutations is smaller still (Fig. 2A).

**Fig. 2.**
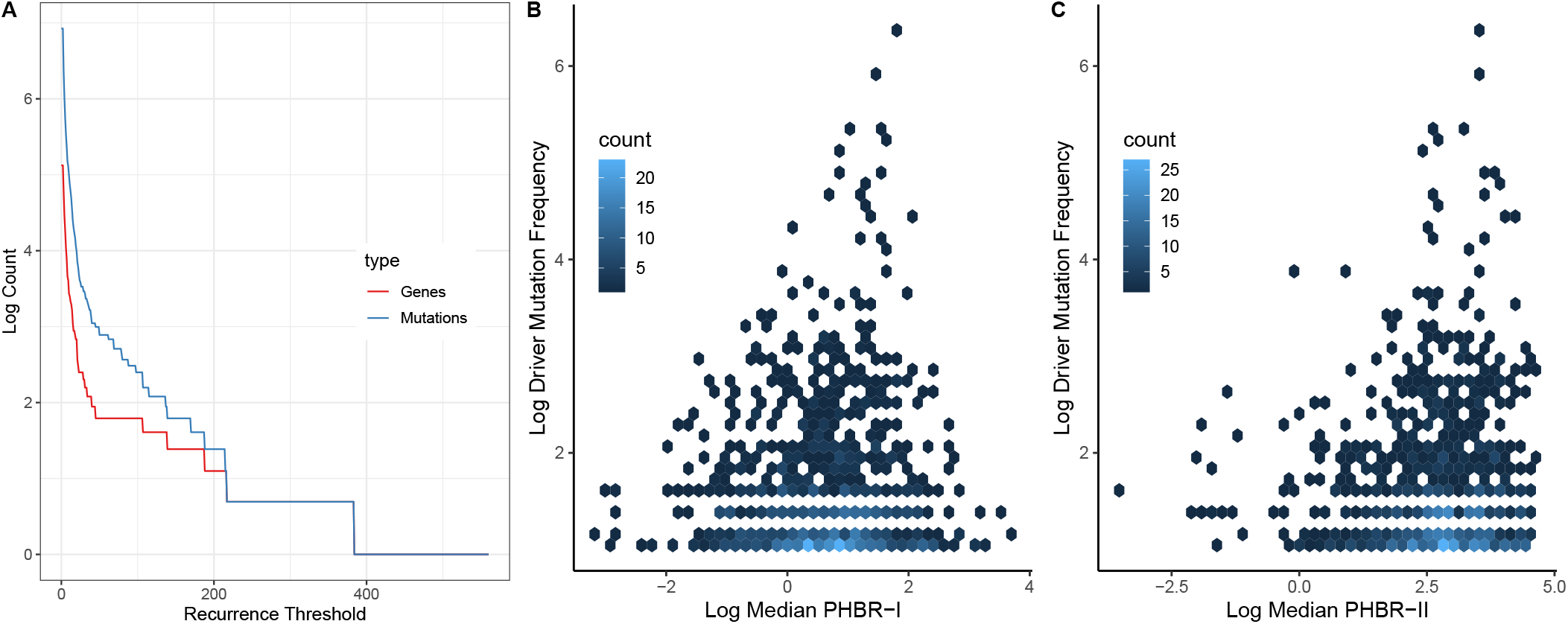
(A) The blue line shows the logarithm of the number of driver mutations that recur across patients at least as often as the recurrence threshold on the x-axis. The red line shows the logarithm of the number of distinct genes in which these mutations occur. (B), (C) Hexbin plots illustrating the relationship between the logarithm of median PHBR-I (B) and PHBR-II (C) scores and driver mutation frequency (across patients).

### Regression models relating log-PHBR score to mutation probability

In addition to comparing PHBR scores between the Mutation and No Mutation groups [16] proposed two mixed effects logistic regression models to relate the log-odds that a driver mutation is found in a patient to the log of the PHBR-I score for the mutation, given patient MHC genotype. In one model (referred to as the within-mutation model) a random effect is used to correct for differences in the frequency with which different driver mutations occur. In the other model (referred to as the within-patient model) the random effect models differences in the abundance of driver mutations between patients, but there is no correction for differences in the frequency of different driver mutations. Mathematical descriptions of both models are reproduced in the Methods.

In [16] there was no significant effect of log PHBR-I on the log-odds of driver mutations using the within-mutation model. Although the results of the within-mutation model are not reported in [17], log PHBR-II is not significantly associated with driver mutation occurrence with this model either. The failure of the within-patient model to detect an effect of log PHBR-I on the probability of a driver mutation was explained in [16] as resulting from the fact that the impact of immune presentation on the probability of a mutation was captured by the random effect. In other words, the tendency for a driver mutation not to be recognized by common HLA alleles resulted in a high driver mutation frequency and this was captured by the random effect in the model. This is not a strong argument, however, because the median PHBR score does not explain much, if any, of the variance in driver mutation frequency in the cancer patients (Fig 2B,C). Even if the variation in driver mutation frequency was entirely driven by MHC class I genotype, it should not fully capture the relationship between driver mutation occurrence and MHC genotype. I.e. the rare driver mutations should still be found associated with the rare MHC genotypes that are not capable of presenting them and the common driver mutations should be found associated with the relatively more common MHC genotypes that cannot present them. This should be detectable with the within-mutation model, even after accounting for differences in driver mutation frequencies.

In contrast to the lack of a signal from the model that accounted for differences in frequencies between driver mutations, Marty and colleagues [16] reported a very strong effect of log PHBR-I on the log odds of driver mutations using the within-patient model (which accounts for differences in driver mutation burden between patients). Quoting a P-value of < 2.2 × 10^−16^, the authors estimate an increase of 28% in the log odds of occurrence of a mutation with each unit increase in log PHBR-I (95% CI: [25%, 31%]). However, this result is affected by the same failure to take account of the non-independence of observations of the same driver mutation that led to the spurious between-group comparisons of PHBR scores discussed above. This can be seen from the fact that the results are not affected by randomization of the patient genotypes. We randomly shuffled the patient genotypes for the real data so that, for each patient, driver mutations were scored with the HLA genotypes of a randomly selected patient. We then fitted the within-patient model to the shuffled data. When we did this we found that the increase in the log odds of a driver mutation occurrence per unit increase in log PHBR-I was 25.1% (standard error 1%), slightly higher than we obtained using the real data (we obtained an estimate of 24.7% when we implemented the within-patient model on the PHBR-I data, a little below the 28% reported by [16]). Similarly, the relationship between PHBR-II was just as strong using the shuffled and unshuffled data (27.0% and 26.9% increase in the log odds of mutation occurrence per unit log PHBR-II for the shuffled and unshuffled data, respectively). Again, these results provide no indication of a relationship between the patient HLA genotypes and driver mutation occurrence.

We performed a simple simulation to demonstrate how the spurious results obtained with the within-patient model can come about. We simulated the case of a single driver mutation that occurs at high frequency and has a high PHBR score across patients. The remaining mutations occurred at lower frequency and had a lower PHBR score distribution (details of the simulation are provided in Methods). Because the within-patient model of [16] and [17] treats PHBR scores of a given mutation as though they were independent observations (despite the strong correlation in the scores of different mutations between patients seen in Figure 1A) this single common driver mutation with a high PHBR score was sufficient to give a highly significant association between PHBR score and driver mutation occurrence (P = 2 x 10^−52^). This trivial example illustrates how failure to account for the high degree of correlation in the immunogenicities of driver mutations across patients can give highly misleading results.

### No evidence that driver mutations in cancer patients are adapted to patient MHC genotypes

Under a null model of no effect of MHC genotype on driver mutation occurrence, the probability that the patient can present a given driver mutation can be estimated from the proportion of all patients that can present that mutation. This provides a straightforward means to compare the observed to expected total number of driver mutations with PHBR scores below the threshold for presentation. If the driver mutation landscape is shaped by patient-specific MHC binding capacity and if this is captured by PHBR scores, then the observed number of driver mutations that can be presented in the patients in which they occur should be smaller than the expected number. For MHC-I the observed number of driver mutations with PHBR-I scores below the threshold of 2 applied in [16] was slightly larger than the expected number (3,669 compared to 3,657.5 ± 68.8). For MHC-II the observed number of driver mutations with PHBR-II scores below the threshold of 10 applied in [17] was slightly below the expected number (1,119 compared to 1,142.3 ± 36.4). Both observed values lay within one standard deviation of the expected values. These results provide no suggestion that driver mutations occur less often in patients with MHC alleles that are capable of binding them.

### Prediction of driver mutation occurrence from MHC genotype

The study of [16] includes the claim that the PHBR scores derived from patient MHC-I genotype could be used to predict the driver mutations that are observed in cancer patients; however, this claim is never tested directly. For each driver mutation, we fitted a logistic regression model to relate the log-odds of a driver mutation occurring as a function of the patient-specific log PHBR-I score. For example, the most common driver mutation in the dataset, V600E in BRAF, occurs in 561 individuals. When we fitted a logistic regression model, treating the log-odds of occurrence of this mutation as the response variable and with log PHBR-I for V600E, cancer type and population of origin of the patient as predictor variables, there was no significant effect of log PHBR-I on the occurrence of this mutation (P = 0.67). It could be argued that common mutations are common because they cannot be presented by common HLA alleles (i.e. they have generally high PHBR scores across patients). While it is the case that V600E in BRAF has a high mean PHBR-I score, there were still 704 patients whose MHC-I alleles were predicted to be capable of presenting this mutation (PHBR-I < 2, the threshold used in [16] to indicate MHC class I binding). Of these patients, 5.5% actually carried the V600E mutation in BRAF, almost identical to the frequency of the mutation in the patients with PHBR-I ≥ 2 (6.2%; P = 0.57 from a Fisher’s exact test). We fitted logistic regression models for each driver mutation and found that no driver mutation was significantly predicted by log PHBR-I, after correction for multiple testing (minimum P value = 0.003; adjusted P = 1, using the Holm method). We repeated this procedure using PHBR-II scores and again found no significant association with driver mutation occurrence following correction for multiple testing (minimum P value = 0.004; adjusted P = 1). There is, therefore, no evidence that patient HLA alleles are predictive of the driver mutations that occur in the patient.

### The association between driver mutation frequency and PHBR scores

The strong associations previously reported between driver mutations and immune presentation scores could be explained by a small number of driver mutations with high frequencies that have high PHBR scores (and therefore are not well presented by HLA alleles). [16] implies that the high frequency of some driver mutations is caused by the fact that these mutations are not well presented by common HLA alleles, thus enabling them to occur in many individuals. This is illustrated by a significant correlation between the frequency of driver mutation occurrence (within bins of driver mutation frequency) and median PHBR-I scores in the bin (this relationship can be seen in the upward trend of the median values from left to right in Figure 1C). Although 1,018 driver mutations were included in the studies of [16] and [17], they are associated with just 168 different genes. Based on an analysis of 1,000 randomly sampled pairs of germline mutations from the same genes, we found that the PHBR scores of mutations in the same gene are positively correlated (Fig. S2), likely reflecting amino acid or domain content of the proteins. For example, peptides of proteins with a large proportion of hydrophobic residues may be more likely to be presented on MHC molecules [20, 15, 21]. The driver mutations with the highest frequencies across patients are dominated by a relatively small number of genes (Fig. 2A). If a subset of these genes tend to have relatively high PHBR scores this could induce a correlation between driver mutation frequency across patients and median PHBR score. Indeed, when we restricted to only the highest frequency driver mutation for each driver gene the relationship between PHBR-I score and driver mutation frequency was no longer significant (Spearman *ρ*= 0.24; P = 0.28). Thus, the reported association between driver mutation frequency and median PHBR-I score is not robust.

### No evidence that driver mutation coverage predicts cancer risk

If the frequency of driver mutations across cancer patients was determined to a large extent by the binding affinities of common HLA alleles, we would expect the number of recurrent cancer driver mutations that can be bound by a patient’s MHC molecules to be associated with cancer risk. In [17] the driver mutation coverage is defined as the number of driver mutations that can be presented by the patient’s MHC molecules. This can be calculated for MHC-I (for which a threshold of PHBR-I < 2 was used to indicate binding) and for MHC-II (for which the threshold was PHBR-II < 10). MHC-I (but not MHC-II) coverage was found to be correlated with age of diagnosis for TCGA patients [16, 17]. Interestingly, the strongest correlations between PHBR-I coverage and age at diagnosis are for cervical and liver cancers, two cancers that are strongly associated with viral infections [22–24], suggesting that the relationship between coverage and age at diagnosis may reflect HLA-dependent differences in susceptibility to these viral infections. To test, more generally, whether there is any relationship between PHBR-I coverage and cancer risk we fitted a logistic regression model to the log-odds of cancer status (a binary variable to indicate whether the individual has self-reported a diagnosis of cancer of any type) to PHBR-I coverage for 377,790 participants from the UK Biobank. Treating age and sex as covariates, we found no significant association between PHBR-I coverage and cancer risk (p = 0.15). The lack of an association between cancer risk and driver mutation coverage does not support a model in which cancer driver mutations occur in gaps in the capacity of the individual’s MHC molecules to bind the associated neoantigens.

## Discussion

The relationship between MHC genotype and the driver mutations that are found in cancer patients, reported by [16] and [17], are unchanged when the MHC genotypes of patients are shuffled. This includes the effect of log PHBR score on the occurrence of a driver mutation, as inferred from the within-patient model, as well as the difference in median PHBR scores between the Mutation and No Mutation groups. It is therefore clear that any effect of PHBR scores on the driver mutation landscape is not dependent on individual level MHC genotypes. It is still conceivable that MHC genotype affects the driver mutation landscape at the population level, such that poorly presented driver mutations are relatively common; however, it is implausible that the population level effect could arise in the absence of any association between PHBR score and driver mutation occurrence within individual patients. If immune responses cause driver mutations that can be recognized by common MHC alleles to be rare, we would expect these driver mutations to be more frequent among individuals with MHC alleles that are incapable of presenting them. No such effect of MHC genotype on driver mutation occurrence within individuals was apparent from the data. Furthermore the relationship that was reported between driver mutation frequency and median PHBR score of the mutation is weak and no longer significant when we restricted to a single driver mutation per driver gene. This restriction is necessary, given the correlation we observed between PHBR scores derived from the same gene, even for germline mutations.

If, as [16] suggests, cancer arises in gaps in an individual’s capacity to present driver mutations, then we would expect the number of such gaps that an individual has for cancer driver mutations to be a strong risk factor for cancer development. Indeed, [17] reports an effect of MHC-I driver mutation coverage on age at cancer diagnosis, where coverage was defined as the number of driver mutations in the study that were predicted to be bound by the patient’s MHC class I molecules. We tested this using data from the UK Biobank. Given the size of the data set (377,790 individuals, including 32,802 with a self-reported cancer diagnosis) even a weak relationship between MHC-I coverage and cancer risk should be detectable; however, we found no significant effect of coverage on cancer status when we fitted a logistic regression model that included sex and age as covariates. If the reported effect of MHC genotype on driver mutation landscape was robust, this would be an important negative result, as it addresses the proposal by [16] that PHBR-I scores of driver mutations may prove useful for assessing risk of development of certain cancers. This negative result has not previously been reported, to the best of our knowledge.

Several studies have reported a depletion of immunogenic nonsynonymous mutations in cancer [13, 9, 14]. However, a recent reanalysis of somatic mutations in cancer that took account of the cancer mutation profiles found no evidence of selection against cancer neoantigens [15], raising questions about whether the availability of neoantigens is the limiting factor in the immune response against cancer. In support of this, a previous study reported that the quantity of neoantigens is not the limiting variable in immunologically cold tumors [25]. This contrasts with studies of the efficacy of immunotherapy which have generally reported a positive association with tumour mutation burden [26– 28]. The reported depletion of cancer neoantigens [13, 9, 14] applies to all nonsynonymous immunogenic mutations and not specifically to driver mutations. However, [16] and [17] reported no evidence of an influence of patient MHC on passenger mutations. This finding is surprising, given that both driver and passenger mutations (particularly clonal, nonsynonymous, immunogenic passenger mutations) should have the capacity to elicit immune responses. In principle, this could be explained by downregulation of genes carrying immunogenic mutations. Indeed, a recent study [29] suggested that the extent of depletion of neoantigens depends on the expression level of the gene. While for neoantigens resulting from passenger mutations, this downregulation might be accomplished without affecting cancer cell proliferation, the requirement of the cancer cells for continued expression of genes carrying driver mutations may prevent downregulation of these genes. One difficulty in attempting to reconcile these findings in this way is that the effect of MHC genotype on the driver mutation landscape was reported for both oncogenes and tumour suppressor genes (and was stronger for the latter group in [17]). It is not clear that the requirement for expression of the gene that carries the driver mutation should apply to driver mutations in tumour suppressor genes, where loss of function is the expected mode of action.

Our reanalysis of cancer driver mutations from the TCGA indicates that there is no evidence that selection exerted by the immune response influences the driver mutations observed in cancer. This result complements the recently reported lack of overall depletion of neoantigens among somatic mutations observed in cancer [15]. It remains possible, however, that the capacity of the MHC to present neoantigens at the cell surface does have an appreciable influence on the driver mutations observed in cancer, but that this capacity is not sufficiently well captured by the PHBR score. Given the experimental evidence for the capacity of PHBR-I and PHBR-II scores to predict MHC-I and MHC-II binding affinity [16, 17], this seems unlikely. Alternatively, it is possible that the availability of immunogenic non-synonymous mutations is not what limits the capacity of the immune response to prevent cancer development. The wide range of mutation burdens in human cancers [30] and the relationship between mutation burden and the efficacy of immune checkpoint inhibitors [26–28] argue against this suggestion, unless the immune response to the developing cancer is distinct to the response following immune checkpoint inhibitor therapy. The lack of a relationship between MHC genotype and driver mutation content suggests that if the immune system plays a major role in cancer prevention this does not involve the prevention of specific driver mutations in a way that depends strongly on MHC genotype.

## Acknowledgements

This publication has emanated from research conducted with the financial support of Science Foundation Ireland under Grant number 16/IA/4612. We acknowledge the QuanTII project under H2020-MSCA-ITN grant agreement number 764698 for support provided to NK. This research has been conducted using the UK Biobank Resource, under Application Number 23739. We thank Rhodri Ceredig for comments on the manuscript.

## Conflict of interest

The authors declare that they have no conflict of interest.

## Notes

### Competing Interest Statement

The authors have declared no competing interest.

