## Supplementary Figures for "No evidence that HLA genotype influences the driver mutations that occur in cancer patients"

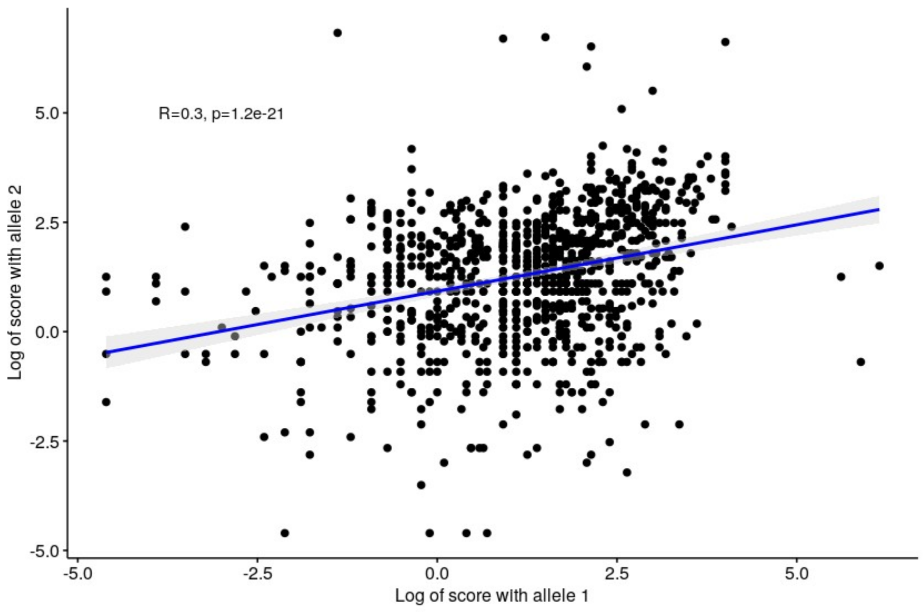


Supplementary Figure 1: Correlations of PHBR-I scores from different HLA alleles

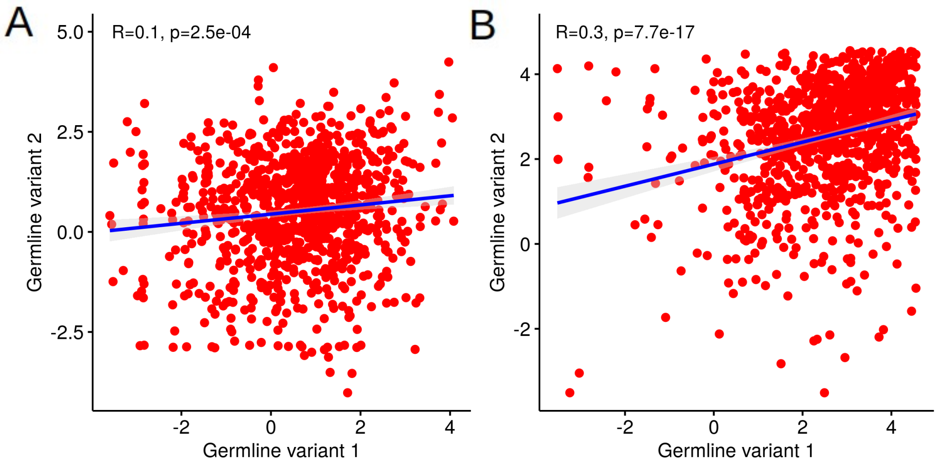


Supplementary Figure 2: Scatterplot of log PHBR-I (A) and log PHBR-II (B) scores for a random set of pairs of germline mutations on the same genes.
